# Bayesian Confidence Intervals for Multiplexed Proteomics Integrate lon-Statistics with Peptide Quantification Concordance

**DOI:** 10.1101/210476

**Authors:** Leonid Peshkin, Meera Gupta, Lillia Ryazanova, Martin Wühr

## Abstract

Multiplexed proteomics has emerged as a powerful tool to measure relative protein expression levels across multiple conditions. The relative protein abundances are inferred by comparing the signal generated by isobaric tags, which encode the samples’ origins. Intuitively, the trust associated with a protein measurement depends on the similarity of ratios from the protein’s peptides and the signal level of these measurements. However, typically only the most likely results are reported without providing confidence for these measurements. Here we present a mathematically rigorous approach that integrates peptide MS-signal and peptide-measurement agreement into an estimation of the true protein ratio and the associated confidence (BACIQ). The main advantages of BACIQ are: 1) it removes the need to threshold reported peptide signal based on an arbitrary cut-off, thereby reporting more measurements from a given experiment; 2) confidence can be assigned without replicates; 3) for repeated experiments BACIQ provides confidence intervals for the union, not the intersection, of quantified proteins; 4) for repeated experiments, BACIQ confidence intervals are more predictive than confidence intervals based on protein measurement agreement. To demonstrate the power of BACIQ we reanalyzed previously published data on subcellular protein movement upon treatment with an Exportin 1 inhibiting drug. We detect ~2x more highly significant movers, down to subcellular localization changes of ~1%. Thus, our method drastically increases the value obtainable from quantitative proteomics experiments helping researchers to interpret their data and prioritize resources. To make our approach easily accessible we distribute it via a Python/Stan package.

## Introduction

Mass spectrometry based proteomics has undergone a remarkable revolution and is now able to identify ~10,000 proteins in a single experiment (1–3). However, due to the difficulty in predicting ionization efficiency of peptides during electrospray, the signal measured in the mass spectrometer is not a direct readout for a peptide concentration in a sample. Proteomics is well suited to comparing the abundance change of the same peptides/proteins among multiple conditions. In so-called label-free proteomics, the peptide signal is compared between multiple different runs and changes of ~2-fold can be detected as significant (4). Even smaller relative protein abundance changes can be detected by encoding multiple conditions with heavy isotopes and analyzing the samples simultaneously. In MS1 based approaches like SILAC the different conditions contain different numbers of heavy isotopes and conditions are encoded by differing peptide masses. However, due to the increase in complexity of the MS1 spectrum with more conditions this approach is only feasible for analyzing up to three conditions at a time (5). A breakthrough for proteomics was the introduction of isobaric tags (6). These tags, which are chemically attached to the peptides, act as barcodes for the different conditions (e.g. replicates, perturbations, or time-points). Each tag has the same mass and only upon fragmentation are the distinct reporter ions released. Due to the identical mass, the MS1 spectrum does not increase in its complexity with more conditions and currently up to 11 conditions can be compared in a single experiment (7). Initially, the co-isolation and co-fragmentation of other peptides led to major artifacts. However, more recently these artifacts have been overcome with the introduction of MultiNotch MS3 (TMT-MS3), QuantMode, and the complement reporter ion approach (TMTc+)(8–12). With these methods, data of superb quality can be generated and abundance changes of ~10% can be quantified as highly significant (13).

Despite these impressive capabilities of quantitative multiplexed proteomics, a remarkable shortcoming is the lack of confidence assigned to these measurements. Typically, only the most likely protein ratios are reported. Various factors can distort the measurements: peptide-to-spectra matching uncertainty, enzymatic digestion efficiency, post-translational modifications, and interference (11). Generally, there is no sense of how much we can trust the data (14). Noise models have been presented to handle peptide-to-protein aggregation in label-free settings. However, these approaches are not easily transferable to multiplexed proteomics, where the data is of a very different nature (15–17). Using multiplexed data, the previous studies considered measurement agreement between peptides assigned to a protein, but the underlying ion-statistics were ignored (18, 19). With replicate experiments confidence can be calculated with standard approaches like the t-test or ANOVA (9, 20, 21). For these approaches, protein level measurements typically weigh peptides by ion-signal but ignore the underlying agreement between peptide measurements. For confidence based on replicate protein-level measurements, the confidence of the measurement can obviously only be expressed for the intersection of protein sets measured in all repeated experiments. Moreover, this approach may lead to unwarranted high confidence when multiple experiments have wrong but concordant measurements and each experiment ignores the disagreement at the peptide level. Also, peptides which are measured with the signal below an arbitrary level are ignored (9, 22).

Most proteins are measured via multiple peptide quantification events. Intuitively, one should be able to use both the agreement between quantifications of peptides assigned to a protein and measured intensities, which are proportional to the number of ions, to assign confidence. Figure S1 summarizes the challenge to express confidence for multiplexed proteomics measurements. However, to our knowledge there is currently no way to integrate all this information to express the confidence of protein level quantification. Assigning confidence is important because it allows one to assess the significance of changes and enables researchers to prioritize valuable time and resources in follow-up experiments.

In this paper, we propose a novel, mathematically rigorous method for computing and representing the uncertainty of quantitative multiplexed proteomics measurements. Our method is based on a generative hierarchical Bayesian model of a two condition Beta Binomial method (or the Dirichlet-Multinomial for the multiple-condition scenario). We call our method BACIQ (**B**ayesian **A**pproach to **C**onfidence **I**ntervals for protein **Q**uantitation – pronounced BASIC). Our approach allows us to represent uncertainty for both individual peptides as well as multi-peptide proteins and considers peptides regardless of the signal level, thereby increasing the sensitivity of proteomics measurements. Our method does not require multiple repeated experiments, but if such repeats are available, it integrates the results providing the output and confidence for the union (not intersection) of all separately measured proteins. If the repeats are available, BACIQ is more predictive than the standard approach using a t-test. Further, we demonstrate the power of our method by re-analyzing a previously published proteomics data-set that studies the change of subcellular protein localization upon addition of the Exportin-1 inhibitor Leptomycin B (13). With BACIQ, we can identify ~2x more Leptomycin B responders compared to our previously published naïve analysis and detect subcellular localization changes of as low as 1% as significant.

## Experimental Procedures

### Sample preparation

The single proteome standard and the two-proteome interference model was prepared mostly as previously described (10, 13, 23). HeLa S3 cells were grown in suspension to 1×10^6^ cells/mL. Cells were harvested by spinning 160 rcf for 5 min at room temperature. After two washes with PBS, the pellet was flash frozen in liquid nitrogen. The pellet containing about 600 μg of total protein was resuspended in 1 ml of lysis buffer containing 25 mM HEPES pH 7.2, 2% SDS and protease inhibitors (complete mini., EDTA-free; Roche). Cells were lysed by sonication: 6 pulses, 10sec each, at 75% amplitude.

*E. coli* cell culture was harvested at 0.5 OD and spun down at 4,000 rcf for 20 min at 4 C. The pellet containing about of 650 ug of total protein was resuspended in 1 ml of lysis buffer containing 8M Urea, 2 M Thiourea, 50 mM HEPES pH 7.2, 2% SDS, 5 mM DTT. Cells were lysed by sonication: 10 pulses, 30 sec each, at 75% amplitude.

200 μL of HeLa lysate was reduced with 5 mM DTT for 20 min at 60 C. Further, both samples – 200μL of HeLa lysate and 200μL of *E.coli* lysate were alkylated with 15 mM *N*-Ethylmaleimide (NEM) for 30 min at room temperature. The excess of NEM was quenched with 5 mM DTT for 10 min at room temperature in both samples. Next, 200 μL of lysate were Methanol-Chloroform precipitated as previously described (24). Protein concentration was determined using the bicinchoninic acid (BCA) protein assay (Thermo Fisher). The samples were resuspended in 6 M Guanidine Chloride in 10 mM EPPS pH 8.5 with a subsequent dilution to 2 M Guanidine Chloride in 10 mM EPPS pH 8.5 for digestion with Lys-C (Wako, Japan) at room temperature with Lys-C 20 ng/μL overnight. Further the samples were diluted to 0.5 mM Guanidine Chloride in 10 mM EPPS pH 8.5 and digested with Lys-C 20 ng/μL, and Trypsin 10 ng/μl at 37 C overnight. The digested samples were dried using a vacuum evaporator at room temperature and taken up in 200 mM EPPS pH 8.0. 10 μL of total *E. coli* or human peptides were labeled with 3μL of TMT 20μg/μL. TMT reagents were dissolved in anhydrous Acetonitrile. TMT Samples were labeled for 2 hours at room temperature. Further, labeled samples were quenched with 0.5% Hydroxylamine solution (Sigma, St. Louis, MO) and acidified with 5% phosphoric acid (pH<2) with subsequent spin at 16,000 RCF for 10 min at 4 C. The samples were dried using a vacuum evaporator at room temperature. Dry samples were taken up in HPLC grade water and stage tipped for desalting (25). The samples were resuspended in 1% formic acid to 1μg/μL and 1μg of each sample was analyzed with the MultiNotch MS3 approach (9).

The samples were labeled with the desired mixing ratios: 1.0:1.0:1.0:1.2:1.2:1.2 for E.coli, and 1.0:1.0:1.0:1.0:1.0:1.0 for HeLa (Figure 6A). Approximately equal amounts of the samples were mixed. To correct for pipetting errors the summed signal for each species was normalized to the desired mixing ratios.

### LC/MS analysis

Approximately 1 μL per sample were analyzed by LC-MS. LC-MS experiments were performed on Orbitrap Fusion Lumos (Thermo Fischer Scientific). The instrument was equipped with Easy– nLC 1200 high pressure liquid chromatography (HPLC) pump (Thermo Fischer Scientific). For each run, peptides were separated on a 100 μm inner diameter microcapillary column, packed first with approximately 0.5 cm of 5-μm BEH C18 packing material (Waters) followed by 30 cm of 1.7-μm BEH C18 (Waters). Separation was achieved by applying 4.8%-24% acetonitrile gradient in 0.125% formic acid and 2% DMSO over 120 min at 350 nL/min at 60C. Electrospray ionization was enabled by applying a voltage of 2.6 kV through a microtee at the inlet of the microcapillary column. The Orbitrap Fusion Lumos was using a MultiNotch-MS3 method (9). The survey scan was performed at resolution of 120k (200m/z) from 350 Thomson (Th) to 1350 Th, followed by the selection of the 10 most intense ions for CID MS2 fragmentation using the quadrupole and a 0.5 Th isolation window. Indeterminate and singly charged, and ions carrying more than six charges were not subjected to MS2 analysis. Ions for MS2 were excluded from further selection for fragmentation for 90 s. MS3 spectra were acquired in the Orbitrap with 120k resolution (200 m/z) simultaneous precursor selection of the five most abundant fragment ions from the MS2 spectrum. The TMTc+ experiments were performed as previously described with 0.4 isolation window on an Orbitrap Lumos (8).

### MS data-analysis

A suite of software tools developed in the Gygi Lab was used to convert mass spectrometric data from the Thermo RAW file to the mzXML format, as well as to correct erroneous assignments of peptide ion charge state and monoisotopic m/z (26). We used ReAdW.exe to convert the raw files into mzXML file format (http://sashimi.svn.sourceforge.net/viewvc/sashimi/). Assignment of MS2 spectra was performed using the SEQUEST algorithm (27) by searching the data against the appropriate proteome reference dataset acquired from UniProt (Escherichia coli (strain K12) – UP000000625 & human) (28) including common contaminants like human keratins and trypsin. This forward database component was followed by a decoy component which included all listed protein sequences in reversed order. Searches were performed using a 20 ppm precursor ion tolerance, where both peptide termini were required to be consistent with Trypsin or LysC specificity, while allowing one missed cleavage. Fragment ion tolerance in the MS2-spectrum was set at 0.02 Th (TMTc) or 1 Th for MutliNotch-MS3. NEM was set as a static modification of cysteine residues (+125.047679 Da), TMT as a static modification of lysine residues and peptides’ N-termini (+229.162932 Da), oxidation of methionine residues (+ 15.99492 Da) as a variable modification. An MS2 spectral assignment false discovery rate of 0.5% was achieved by applying the target decoy database search strategy (29). Filtering was performed using a linear discrimination analysis with the following features: SEQUEST parameters XCorr and unique Δ XCorr, absolute peptide ion mass accuracy, peptide length, and charge state. Forward peptides within three standard deviation of the theoretical m/z of the precursor were used as positive training set. All reverse peptides were used as negative training set. Linear discrimination scores were used to sort peptides with at least seven residues and to filter with the desired cutoff. Furthermore, we performed a filtering step towards on the protein level by the “picked” protein FDR approach (30). Protein redundancy was removed by assigning peptides to the minimal number of proteins which can explain all observed peptides, with above described filtering criteria (31, 32). We only used quantitative information from MS3 and low m/z MS2 reporter ions if we observed an isolation specificity of at least 75% in the MS2 isolation window. We did not use isolation specificity filtering for the TMTc+ method, as co-isolation of other peptides does not perturb the measurement results for this method (8). TMTc+ data were analyzed as previously described (8). The mass spectrometry proteomics data have been deposited to the ProteomeXchange Consortium via the PRIDE (33) partner repository with the dataset identifier PXD012285 with username and password 3I66zA6u.

### Implementation of BACIQ

We develop a hierarchical beta binomial model to assign confidence to the estimate of protein ratio in different conditions (Fig. 5). The model is reparametrized to make the true protein ratio as a parameter. The parameters for the beta distribution were modified into μ,κ from α,β; where 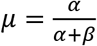 and κ = α + β. The prior for the true protein ratio μ is set to follow a uniform distribution; which indicates that we do not have any prior knowledge for the true protein ratio. The histogram of the samples for μ represents the marginal posterior distribution of the true protein ratio conditioned on the observed data, and hence gives the uncertainty associated with it. The model and the inference was implemented using Stan (34), specifically the Python flavor. Stan’s sampling functionality was used to execute Monte-Carlo Markov Chain to obtain a sample from the posterior distribution over parameters of the model. Since for any values of the model’s parameters we can efficiently evaluate the likelihood, parameter space is explored while samples are biased towards the areas of high likelihood. The obtained statistical sample is essentially used for a histogram representation of the posterior to obtain confidence intervals.

## Results

### Peptide measurements map to coin-flips

To reason about measuring confidence in multiplexed proteomics, let us begin by discussing the measurement of peptides that are labeled with isobaric tags encoding two different conditions (e.g. case and control) (Fig. 1A). With multiplexed proteomics we can measure the relative abundance of peptides between multiple conditions. For the sake of simplicity, we will discuss the two-condition case for most of the paper, but all our approaches and the provided code can be generalized to the multi-condition case. Once peptides from different conditions are labeled with isobaric tags, they are combined into one test tube, in which we have a “true ratio” of peptide abundances across conditions. The aim of the proteomic experiment then is to recover this “true peptide ratio” and assign confidence to the measurement. During MS analysis, the peptides get ionized and fragmented. Upon fragmentation, respective fragments of the isobaric tag are released, encoding the different conditions. The relative abundance of the ions encoding the different conditions is used to quantify the relative abundance of the peptides in the two conditions (Fig. 1B). However, the limited number of ions by which measurements are performed introduce measurement errors due to ion-statistics. The process of estimating the “true peptide ratio” and assigning confidence to the estimate is analogous to evaluating the confidence in fairness of coin θ, given the number of heads and tails (α, β) in n coin tosses. Using the Bayesian approach this process can be described by the standard Beta distribution (Fig. 1C) (35). Intuitively, the confidence interval tends to be wider for measurements with lower signal, which is proportional to the ion-count (Fig. 1D). Isobaric tags are most commonly read out in the Orbitrap mass analyzer, where the raw signal divided by the Fourier transform-noise is proportional to the ion-count (36). Throughout the paper we will refer to this read out as “MS-signal”. To be able to apply the Beta distribution to the proteomics data, the challenge is to find the proportionality constant for converting MS-signal into the number of ions.

**Figure 1.**
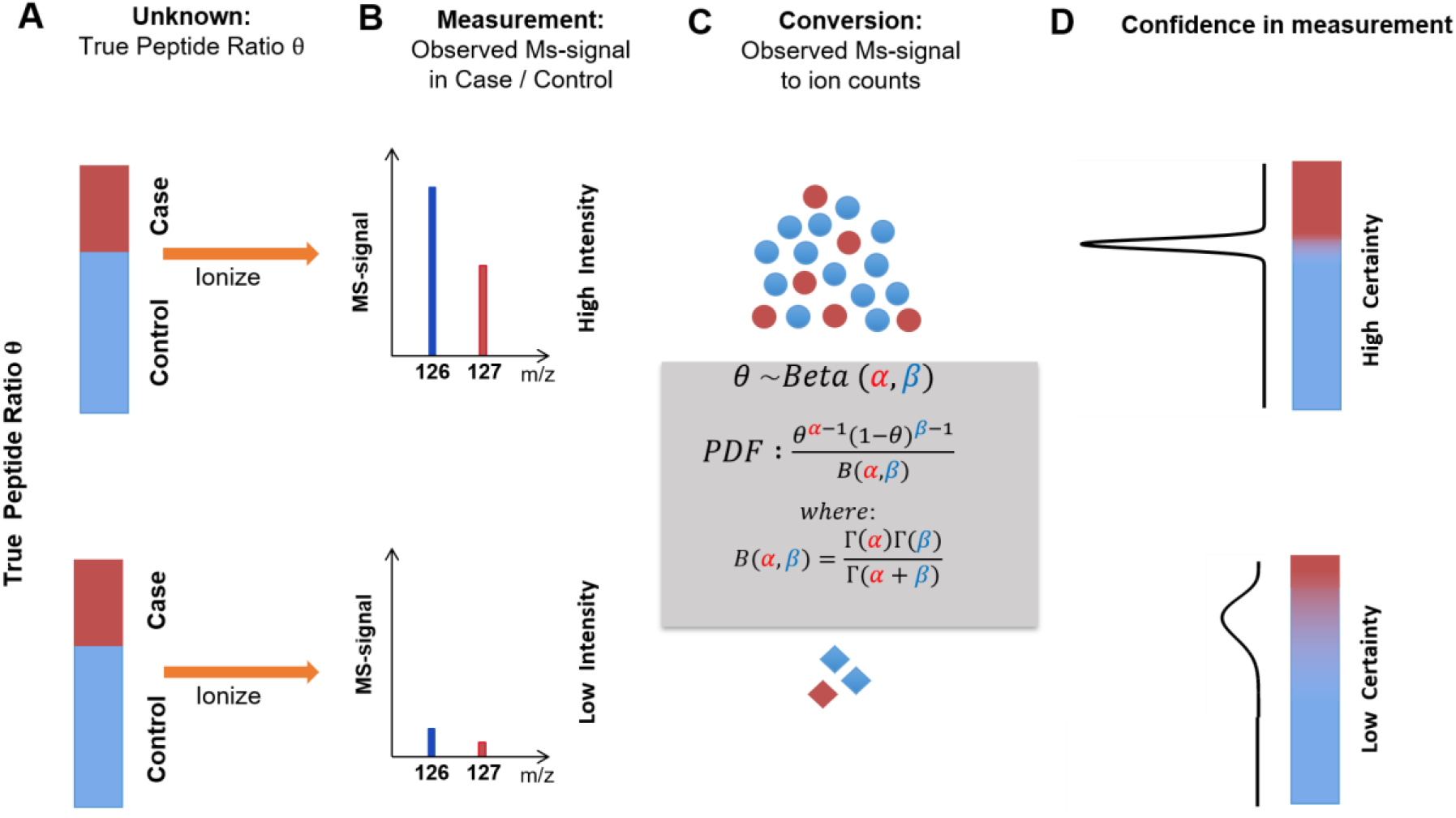
Peptide measurements are analogous to coin-flips: A) Peptides are labeled with isobaric tags encoding the different conditions case and control (red and blue). Shown are examples for two peptides with the identical relative abundance (true ratio) between case and control. B) After ionization and fragmentation, the relative intensity of fragments produced by the isobaric tag can be used to quantify relative peptide abundance. These quantification spectra not only contain information about the peptide relative abundance but also the MS-signal. This signal is proportional to the number of ions measured. C) The problem of estimating the likelihood of the “true ratio” becomes identical to the estimation of a coin’s fairness, given a certain number of head and tail measurements. Using a standard Bayesian approach, we can express the underlying “true ratio” likelihood as a beta distribution where *α, β* represent the number of ions measured for the two samples. D) Intuitively, the fewer ions we measure the more the measured ion ratio tends to divert from the true ratio between case and control due to ion-statistics. A higher ion-count (top row) results in a tighter likelihood function and respectively tighter confidence intervals than a low ion-count (bottom row). For more than two cases this approach can be generalized with a Dirichlet distribution.

### Conversion of mass spectrometer signal to ions

The peptide ratio measurement converges to a true ratio when measured with high MS-signal. To deduce how MS-signal relates to confidence, we generated and analyzed a proteomics sample in which all peptides are labeled with the identical 1:1 ratio. To this end, we mixed the TMT-NHS ester solutions with the desired ratio before adding peptides for labeling. In this experiment, measurement distortions are predominantly due to ion statistics. Upstream sources of noise e.g. differential labeling or digestion are eliminated. The peptides were analyzed with TMT-MS3 or via the complement reporter ion approach (TMTc+) on an Orbitrap Lumos (8, 9). Figure 2A presents a scatter plot where each point represents a single peptide from a TMT-MS3 dataset of a total of ~10k quantified spectra. We observe that the fraction of the signal converges to an asymptotic value, just like the fractional outcome of a sequence of coin-tosses converges to a true fraction with the increasing number of tosses.

**Figure 2.**
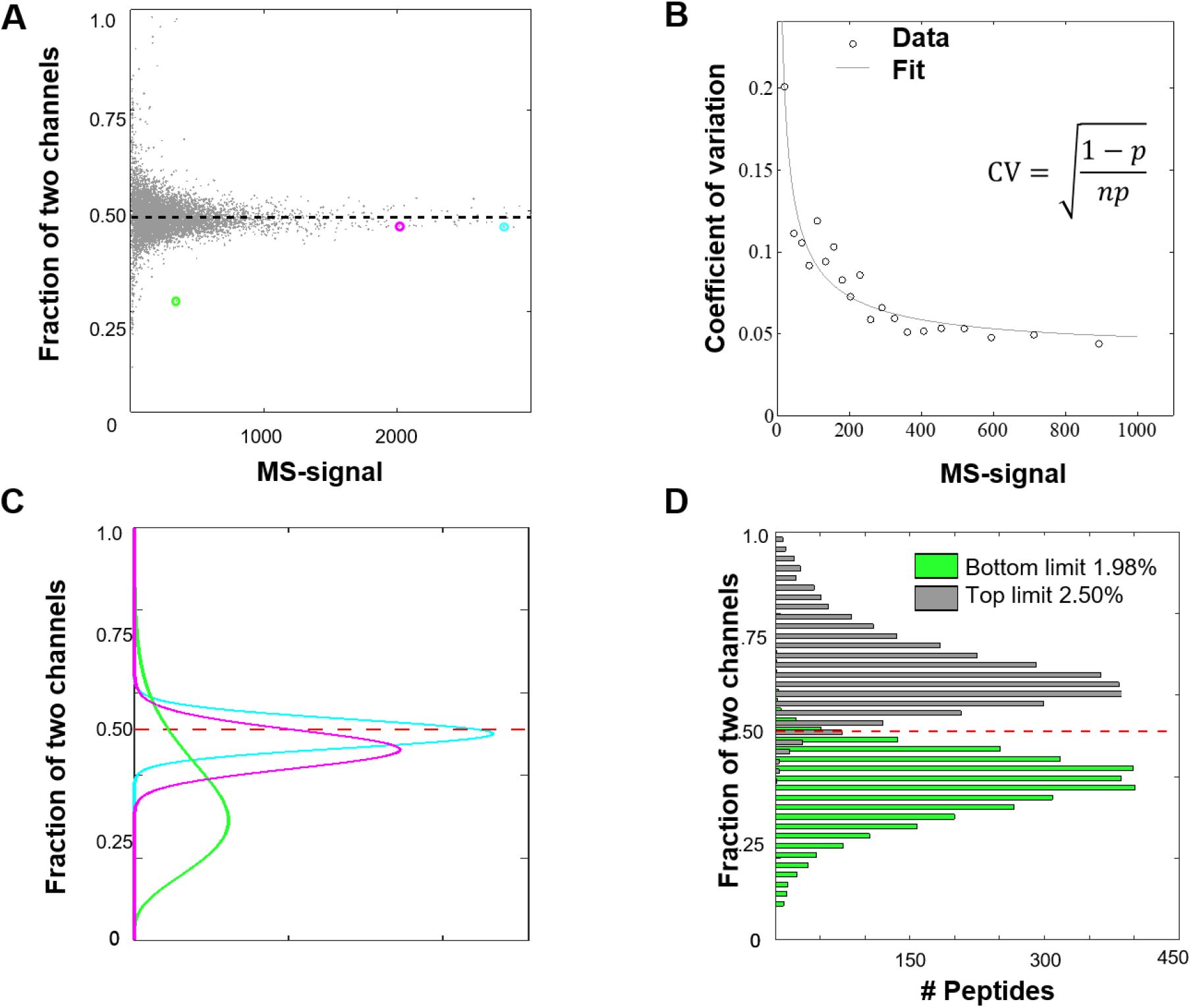
Conversion of MS-signal into counts and assigning confidence to the measurement of a single peptide. A) We generated a sample in which all peptides are labeled with two different TMT-tags and mixed with identical ~1:1 ratio. When we plot the measured fraction in one channel versus the summed MS-signal in both channels, we observe that measurement with higher MS-signal asymptotes to true underlying ratio (dashed line). B) Assuming ion-statistics is the dominant noise source, we can fit the coefficients of variation (CVs) and obtain the conversion factor of MS-signal to the number of ions or pseudo-counts. The data shown was obtained on an Orbitrap Lumos with 50K mass resolution. Our best estimate for the conversion factor is 2.0. C) Three examples color-coded to correspond to three peptide data points in sub-figure (A), of the likelihood functions reflecting confidence, expressed as the beta-distribution using the calculated ion-counts. D) Histogram of the upper and lower bound values for the 95% confidence intervals. The observed percentage of peptides for which the true answer is outside of the 95% confidence interval is 1.98% and 2.5% respectively for over- and underestimation, which are symmetric and in good agreement with the expected total 5%.

This functional form of the convergence of the true fraction of successes with n coin tosses is represented with coefficient of variation of a Binomial distribution 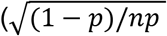, where ***p*** is the fairness of the coin and n the number of tosses). To treat the continuous MS-signal as an equivalent to the number of coin flips, we fit a single parameter ***m*** as a multiplier to an MS-signal s where ***n = ms*** to the data binned on the MS-signal (Fig. 2B). When we perform this analysis on an Orbitrap Lumos for a TMT-MS3 experiment with 50K mass resolution, we observe a conversion factor of 2.0. Interestingly, the conversion factor differs when we repeat the equivalent experiment on a different instrument (Orbitrap Elite), with different resolutions, or methods for data acquisition (Table 1). Makarov et al. previously reported that this conversion factor should scale inversely with the square-root of the Orbitrap resolution (36). Our measurements are in rough agreement with this prediction; when we increase resolution from 15K to 120K on the Lumos we expect the conversion factor to reduce by 2.8-fold, we observe a 3.5 decrease. We suspect that the different conversion factors on different instruments is due to differences in Orbitrap electronics and data processing. For a limited number of cases, we have repeated these measurements for various instruments of the same model and obtained very similar results suggesting that for a given instrument model and resolution the conversion factors are invariant. We observe slightly smaller conversion factors for TMTc+ data into apparent counts compared to TMT-MS3 data. This is most likely due to some additional noise that is introduced during the deconvolution process of the complement reporter ion clusters (8, 10). Based on previous reports (36), we likely underestimate the number of actual ions by a small factor. Nevertheless, the good fit to the data (Fig. 2B) indicates that the conversion into pseudo-counts allows us to model the relationship between mass spectrometer signal and measurement noise due to ion-statistics. Importantly, this calibration step has to be performed only once for any type of instrument and is conveniently supplied here for two commonly used mass spectrometers at various resolutions (Table 1).

**Table 1.** Multiplier for the MS-signal to be converted into discrete Binomial event counts. Shown are the multiplier values and respective confidence intervals for the Orbitrap Lumos or Orbitrap Elite for TMT-MS3 or TMTc+ experiments and various Orbitrap resolutions. *extrapolated from the other values.

| **Instrument** | **Resolution** | **Multiplier low**<br>**m/z TMT** | **Multiplier TMTc+**<br>**(0.4 Th)** |
| --- | --- | --- | --- |
| **Orbitrap Elite** | **15K** | **4.5** | **N/A** |
| **Orbitrap Elite** | **30K** | **3.3** | **N/A** |
| **Orbitrap Elite** | **60K** | **2.5** | **N/A** |
| **Fusion/Lumos** | **15K** | **3.4** | **2.7\*** |
| **Fusion/Lumos** | **30K** | **2.6** | **2.1** |
| **Fusion/Lumos** | **50K** | **2.0** | **1.9** |
| **Fusion/Lumos** | **60K** | **1.8** | **1.7** |
| **Fusion/Lumos** | **120K** | **1.3** | **1.3** |

### Assigning confidence intervals for individual peptides

With the conversion factor at hand, we can convert the MS-signal into the number of observed ions and approximate the confidence associated with the fraction of peptide with a Beta distribution. Figure 2C illustrates the Beta likelihood function of the probability of fraction for three peptides at the various levels of total MS-signal as color-coded in Figure 2A. A higher signal gives a tighter distribution.

We next verify that the confidence intervals obtained agree with observations. Computing the 95% confidence intervals we expect that the true answer will lie outside of the confidence interval approximately 5% of the time and will be symmetrically split between over- and underestimation. Indeed, Figure 2D shows that we overestimate 1.98% of the time and underestimate for 2.50% of the time. We repeated this demonstration for peptides labeled in different ratio and obtained consistent results, showing that beta distribution indeed is a good general model for expressing confidence for single peptide measurements (Fig. S2).

### Only considering ion-statistics produces inadequate confidence intervals on the protein level

So far, we have shown that we can adequately express the confidence intervals for peptide measurements. If ion-statistics was the only source of noise, we could sum up all counts from peptides mapped to a particular protein and express confidence intervals at the protein level. This approach works well for the synthetic experiment, where all peptides in a mixture were labeled together and show the exact same ratio (Fig. S3). However, in real experiments, other factors like differences in digestion efficiency, labeling problems, erroneous peptide-to-protein assignment, post-translational modifications, chemical interference, and so forth might produce significant additional noise. To test whether only considering ion-statistics is valid for multiplexed proteomics measurements, we revisited our previous publication of nucleocytoplasmic partitioning in the frog oocyte (13) (Fig. 3A). We observe that the **R**elative **N**uclear Concentration (RNC) [Nuc/(Nuc+Cyto)] measurements for peptides for one protein disagree with each other. This is shown by the mutual exclusiveness of the confidence interval for the two extreme peptides of one protein (Fig. 3B). Moreover, when we evaluate the confidence based on the sum of all the peptides mapped to a protein, we observe that their probability distribution is unjustifiably narrow (Fig. 3B). This suggests that besides ion-statistics, other significant sources of error contribute to the errors in proteomics measurements and we must take these sources into account to adequately express confidence intervals at the protein level.

**Figure 3.**
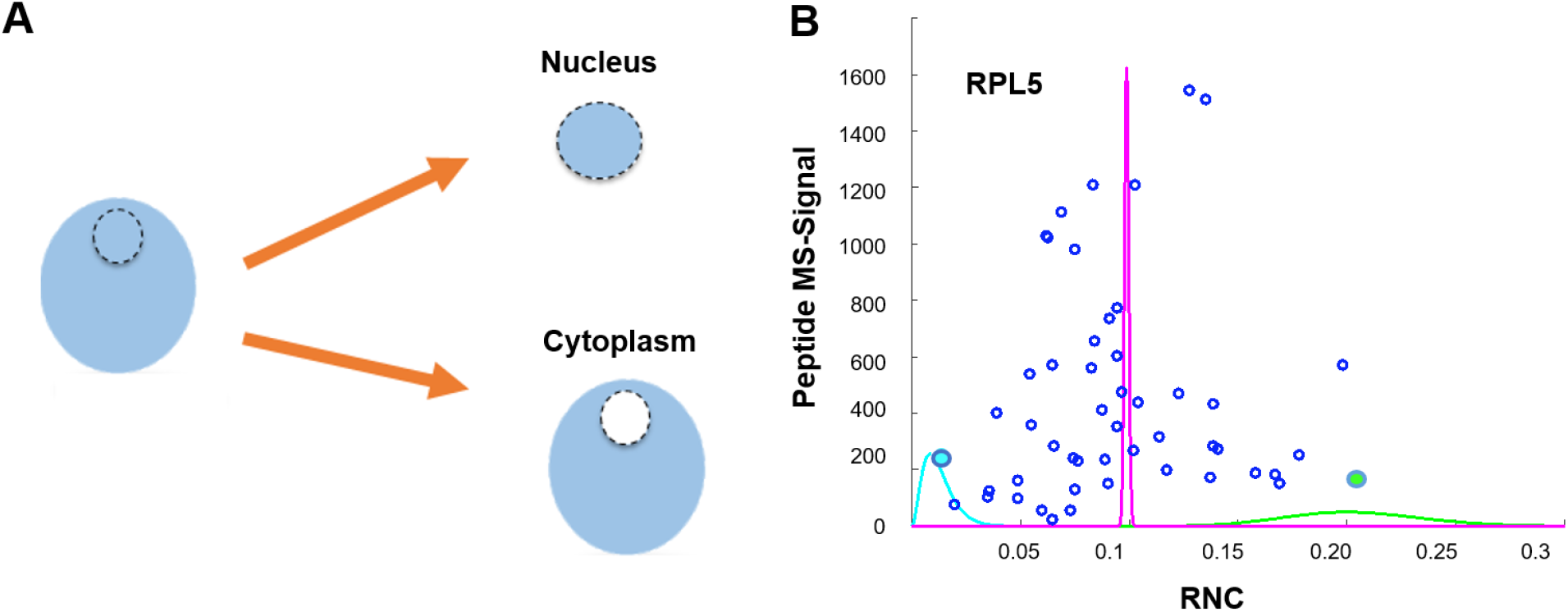
Only considering ion-statistics does not produce accurate confidence intervals at the protein level: A) To evaluate the confidence intervals of peptides from the same protein, we revisited our previously published experiment, where we measured the localization of proteins between nucleus and cytoplasm in the frog oocyte. B) Blue discs show 50 measured peptides (**R**elative **N**uclear **C**oncentration (RNC) [Nuc/(Nuc+Cyto)] and MS-signal) assigned to the Ribosomal Protein L5 (RPL5). We show the beta-function likelihoods for two extreme peptides (leftmost in blue and rightmost in green). Note that these peptides’ likelihood functions are basically mutually exclusive, i.e. the most generous confidence intervals would exclude each other. Additionally, we show the likelihood based on summing up all the peptides together (magenta) which corresponds to unjustifiably tight confidence. This example illustrates that for the expression of confidence intervals on the protein level, we cannot assume that ion-statistics is the only source of measurement error in proteomics experiments. Rather, we have to integrate other sources of errors e.g. due to differences in sample handling.

### Confidence-intervals at the protein level, that integrate ion-statistics and agreement between peptides mapped to the same protein

The goal is to develop a model that gives out a probabilistic distribution for the true fraction in a channel for every protein. Mathematically, achieving the above objective involves calculating the conditional probability of a true protein fraction given the observed individual peptide data – called the posterior distribution. Bayes theorem lets us model this distribution by asking the same question in a converse, more tractable manner i.e. what is the likelihood of observing the peptide measurement data for different protein fractions? This likelihood function can be approximated by reviewing the entire proteomics experiment as a data generating process, starting with a true protein ratio (Fig. 4). The protein is digested into peptides, which are labeled with isobaric tags. This process can introduce disagreement between different “true peptide ratios” because of the differences in sample handling e.g. differing digestion or labeling efficiencies between case and control. We represent this first step as an equivalent of sampling peptide ratios for the constituent peptides of a given protein from the probability distribution parameterized according to the “true protein ratio” (Fig. 4A). In the second step the peptide is ionized and fragmented and its ratio is measured on the mass spectrometer. Due to the limited number of ions measured for each peptide, the observed fraction deviates from the true peptide fraction. This step can be represented as sampling the observed data from a probability distribution parameterized over true peptide ratio, for each peptide separately. Importantly, different peptides are measured with different MS-signal and therefore with different confidence on the underlying “true peptide ratio” (Fig. 4B). We have adequately addressed this second step above. However, we aim for an approach that integrates ion statistics and agreement between peptide measurements to estimate the true protein ratio and the confidence associated with it (Fig. 4C).

**Figure 4.**
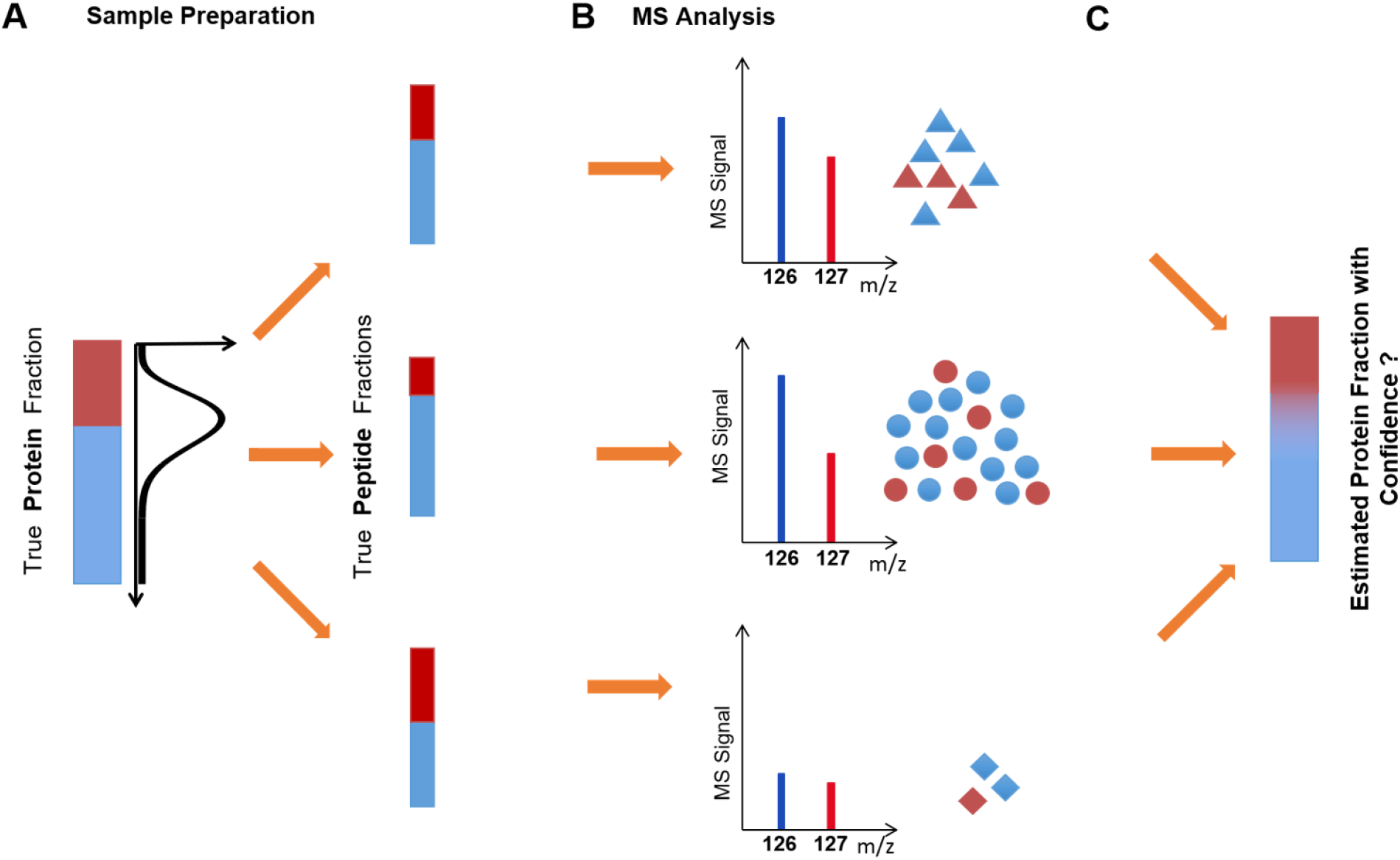
Schematic of the data generating process for modeling confidence for proteins with multiple peptide measurements. A) The “true protein ratio” can be distorted due to differences in sample handling (e.g. digestion, isobaric-labeling, post-translational modifications) and give rise to multiple peptides with differing “true peptide ratios”. The peptide ratios for the constituent peptides of a given protein are sampled from the probability distribution parameterized according to the “true protein ratio” B) Each peptide is measured via the mass spectrometer. Based on the number of ions used to measure each peptide, the confidence in quantification varies. The observed data is sampled from a probability distribution given a true peptide ratio, for each peptide separately C) The goal is to infer the underlying true protein ratio between the conditions and generate confidence using the agreement between multiple peptide measurements and their respective MS-signals.

This entire description of the process can be mathematically represented as a generative two-level hierarchical Beta-Binomial model (35). True peptide ratios *θ*_1_, *θ*_2_,… *θ_i_* are sampled from a beta distribution with parameters *α,β* (***θ_i_** ~ **Beta** (**α,β**)*). Given true underlying peptide ratios, the number of ions in Case channel *y*_1_, *y*_2_,… *y_i_* can be modelled using a Binomial distribution **(*y_i_* ~ ***Bin***(*n_i_, θ_i_*), *where n_i_ is the total number of ions*)** (Fig. 5). Because no other information, other than the observed peptide data y is available, all the unobserved parameters (*α,β,θ*_1_, *θ*_2_,… *θ_i_*) are modeled by a joint probability distribution conditioned on the observed data y. Since each of the peptide measurements are independent from each other, the probability distributions for every peptide can be modeled separately and multiplied together (Fig. 5).

**Figure 5.**
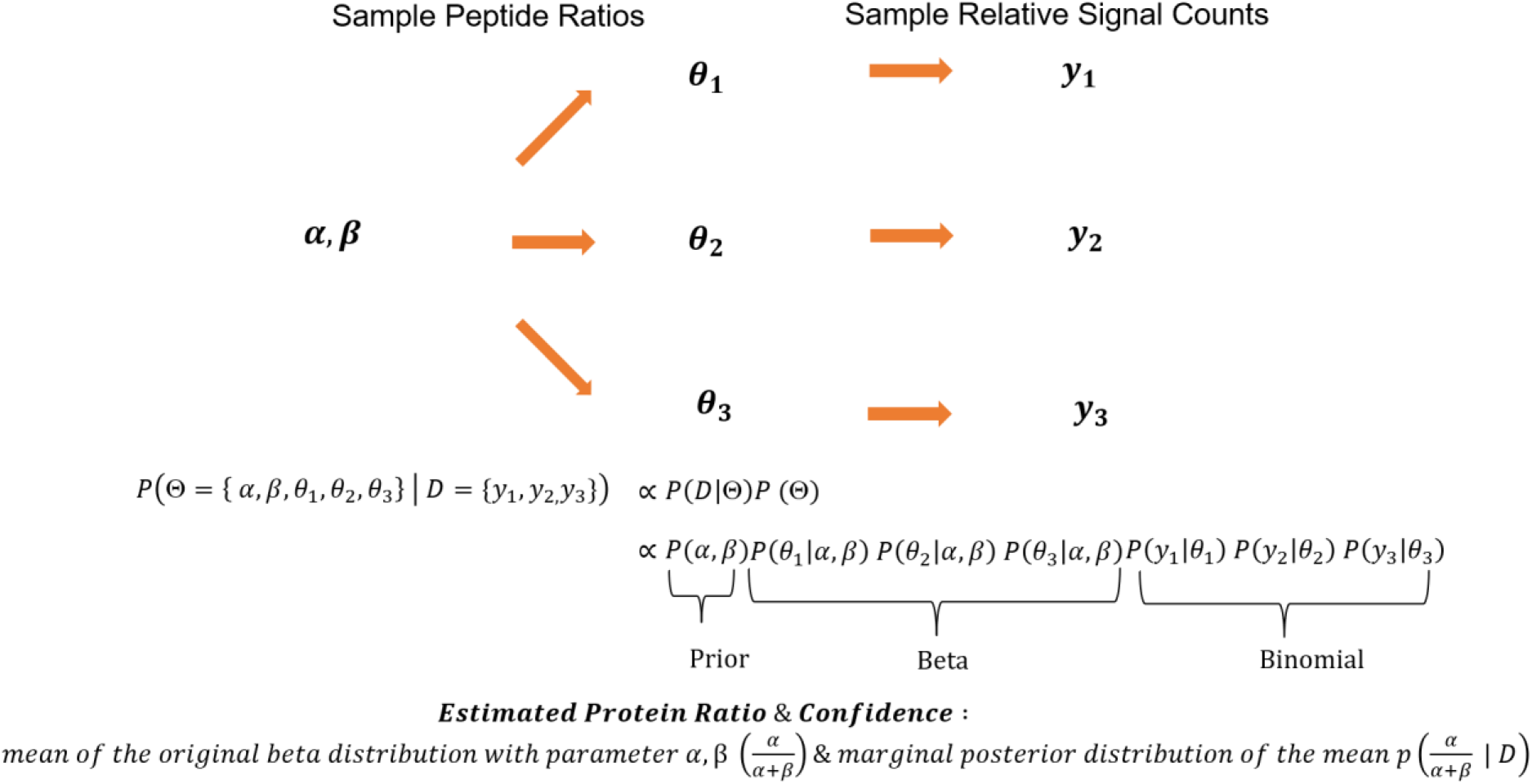
Mathematical model for estimating the protein ratio and its confidence: The entire data generation process can be adequately described with the Beta-Binomial process (or Dirichlet-Multinomial process for more than two cases). We assume an underlying beta-distribution (*α,β*) representing the likelihood function over true protein ratio, from which peptide ratios for the constituent peptides of a given protein is sampled, for i=1, 2 and 3. Given a true underlying peptide ratio and observed number of ions in red channel (*y_i_*), we can sample from respective Binomial distribution with probability of success *θ_i_*. We can use this mapping and express confidence as an estimation for the mean of the Beta distribution representing the protein, specifically the probability of marginal posterior distribution of the mean as the representation of the uncertainty.

Naturally, the target of our estimation (true protein ratio) is a function of the parameters (*α,β*) of original beta distribution. We assume it to be the mean of the beta distribution given as 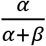 and the uncertainty associated with this estimate is represented by the marginal posterior distribution of this mean conditioned on the data y.

The maximum-likelihood estimate (MLE) of these distributions is not available in a closed form, thereby making the problem analytically intractable (37). One approach would be to numerically search for an MLE and estimate the curvature of the likelihood function from the Hessian at the MLE using the asymptotic normality (38). Unfortunately, however, this approach did not turn out to be numerically robust (not shown). A robust alternative approach is accomplished using Monte-Carlo Markov Chain (MCMC) methods implemented in statistical inference language Stan (see Supporting Information)(34). Essentially, it consists of exploring the space of possible protein ratios, computing the likelihood of observed peptide data given a guess at the protein ratio and assembling a large set of plausible ratio samples to use its histogram as posterior likelihood representation. Mathematical details of the model implementation and reparameterization to obtain samples for true protein ratio are discussed in the Materials and Methods section. The source code for BACIQ in the Python flavors is available via our website: https://scholar.princeton.edu/wuehr/tools-and-resources/BACIQ

### Verification of protein level confidence interval estimation

To verify the power of our approach for detecting significant protein abundance changes, we compared it to the current standard approach of using t-tests on replicate measurements. To this end, we produced a standard containing peptides with different labeling ratios. We asked how reliable our approach was in distinguishing proteins which show different expression levels (increase by 20%) and proteins that are unchanged. We mixed equal amounts of human proteins across six samples with *E. coli* proteins in different mixing ratios in triplicates (Fig. 6A). Thus, ideally all *E. coli* proteins should be identified as “differentially expressed” on a background of uniformly expressed human proteins. This synthetic experiment simulates an essential application of having the complete likelihood function, where confidence intervals representation is used to prioritize the follow-up targets of a proteomic experiment. Crucially, while the t-test requires at least one repeat, our method can be applied to a single experiment, as well as two and three repeats by merely combining measurements. As shown in Figure 6B, even using no replicates our method outperforms the t-test based classification with two repeats for most false positive (FP) rates. For the same number of replicates, BACIQ outperforms the t-test for all FP-rates. Using three replicates, BACIQ achieves a close to perfect distinction. Similar outperformance of BACIQ over the t-test can also be observed for 1.1-fold and 1.4-fold changes (Fig. S4).

**Figure 6.**
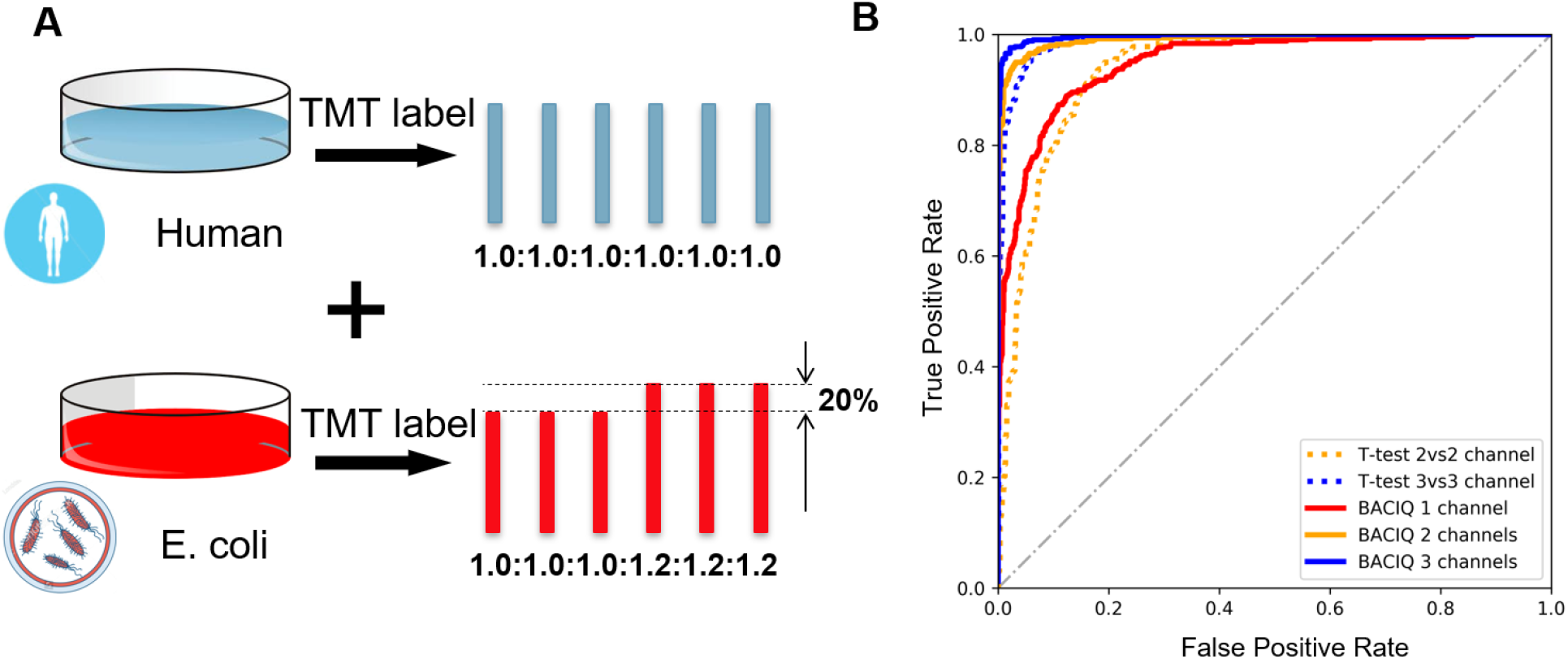
Validating our method with a differential expression experiment. A) Six samples were prepared by mixing material from two species as follows. Six identical human samples (i.e. proportions across 6 channels were 1.0: 1.0: 1.0: 1.0: 1.0: 1.0) were mixed with an *E. coli* sample in two sets of three as shown (i.e. proportions across 6 channels were 1.0: 1.0: 1.0: 1.2: 1.2: 1.2). A mixture of peptides from the two proteomes was analyzed by LC-MS. B) Comparison of our method with a one tailed t-test to detect significantly changing proteins. BACIQ can detect statistically significant changes with a single comparison, while the t-test requires replicates. ROC plot indicates that our method is superior to the t-test when the same number of replicates are used. Even without replicates our method (red) nearly outperforms the t-test with two replicates (dashed yellow). We achieve close to perfect detection of the significantly changing proteins by using the BACIQ analysis with three replicates (blue).

### Re-analyzing subcellular re-localization data with BACIQ increases detection of significant responders to Leptomycin B by ~2-fold

To assess BACIQ under real-world scenarios, we reanalyzed a previously published proteomic dataset that investigated the change of subcellular localization upon Leptomycin B (LMB) addition (13). Leptomycin B is a highly potent and specific inhibitor of Exportin-1 (39), which is responsible for the transport of proteins out of the nucleus into the cytoplasm. Upon treatment, we expect substrates of Exportin-1 to move towards the nucleus (Fig. 7A), due to competing transport with importins or due to passive diffusion through the nuclear pore. Importantly, in this dataset, we can use hundreds of proteins small enough to diffuse through the nuclear pore to normalize for equivalent amounts of nuclear and cytoplasmic material (Fig. S5).

**Figure 7.**
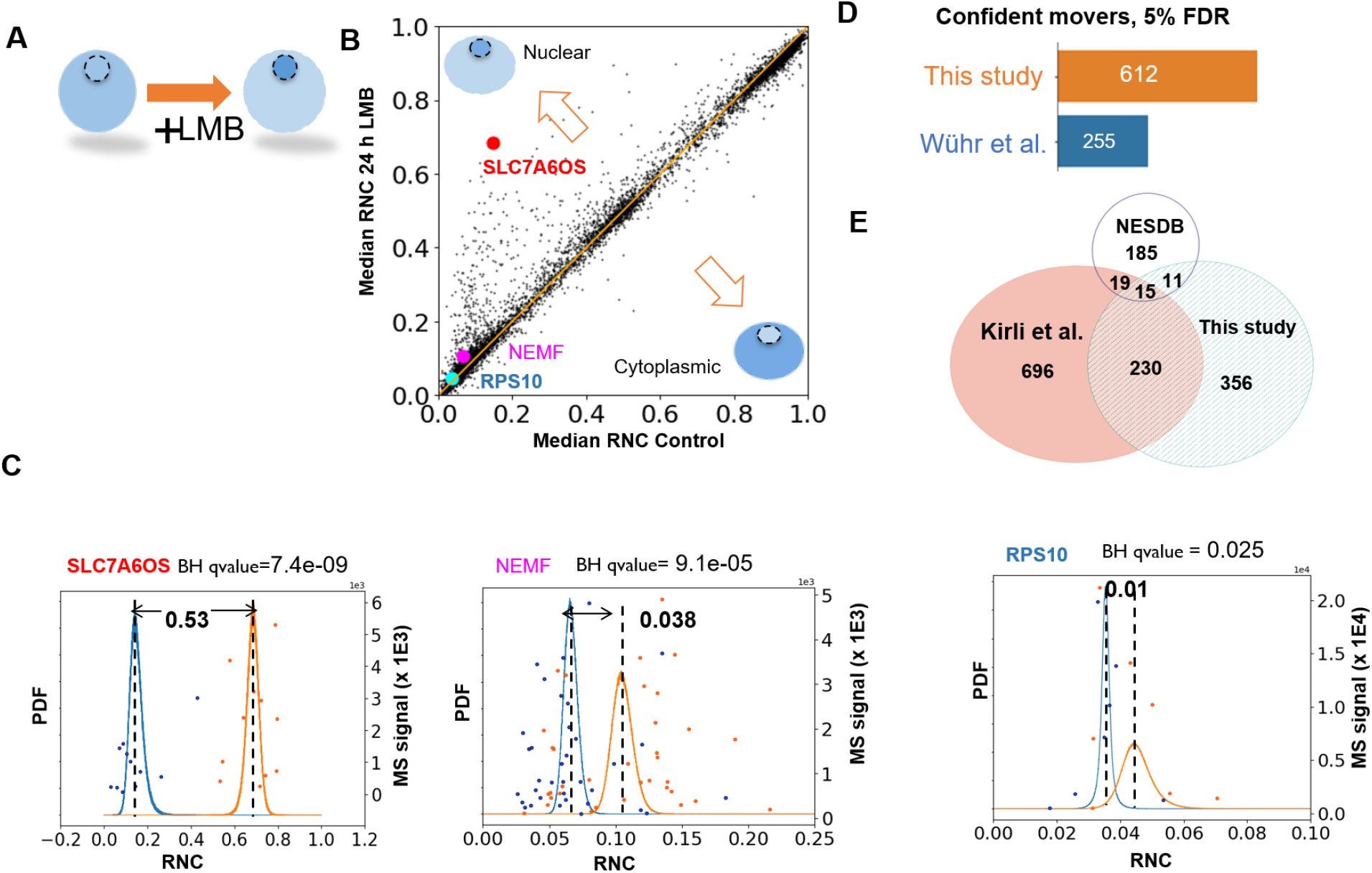
Re-analysis of subcellular movement upon Exportin-1 inhibition with BACIQ. A) Upon inhibition of Exportin-1 with Leptomycin B (LMB) we expect Exportin-1 substrates to move towards the nucleus. We compared the **R**elative **N**uclear **C**oncentration (RNC) [Nuc/(Nuc+Cyto)] in control and drug treated samples to identify the proteins that confidently shift towards the nucleus regardless of initial nucleocytoplasmic distribution. B) A scatterplot indicating the shift in RNC post LMB treatment. Most proteins seem unaffected by the treatment. The proteins above the diagonal indicate movement towards the nucleus and those below the diagonal indicate movement towards the cytoplasm. C) The raw peptide data and probability distributions of RNC for three different proteins (shown as discs in B). The blue curve represents the probability distribution of RNC before adding the drug and the orange curve represents the probability distribution of RNC after adding the drug. With our approach, we can detect the movement towards the nucleus of as little as 1% D) Applied to the entire data-set BACIQ detects 750 putative Exportin-1 substrates (612 unique gene symbols) at 5% false discovery rate. With identical FDR, BACIQ extracts ~2x more proteins as significantly moving compared to our previously published naive analysis. E) Venn diagram shows the overlap in unique gene symbols of the LMB responders at 5% FDR from this study with Cargo database based on Exportin-1 affinity experiments (Kirli et al.), and a database curated from literature (NESDB). The overlap with both databases is highly significant with p values of 5.5 e-29 (Kirli et al.) and 3.1e-6 (NESDB) based on Fisher’s exact test.

Upon comparing the median **R**elative **N**uclear Concentration (RNC) [Nuc/(Nuc+Cyto)] for control and drug treated samples, we observe that most of the proteins do not show any obvious shift. Proteins off the diagonal show apparent movement towards the nucleus or cytoplasm (Fig. 7B). Fig. 7C shows RNC distributions from the Beta-Binomial model of the control (blue) and drug treated (orange) samples for three different proteins. Naively, as we did in our previous publication, one could rank proteins by the magnitude of their movement towards the nucleus as likely Exportin-1 substrates. BACIQ, however, not only allows us to consider the magnitude of movement but assign probability to the movement using the RNC distributions of control and drug treated samples.

In large scale datasets the probability assigned to a single discovery is not particularly meaningful as thousands of hypotheses are tested. We must therefore apply a multiple hypotheses correction procedure. In the discussed experiment, we can use movement towards the cytoplasm upon LMB treatment as a conservative noise model. Indeed, we observe that the p-values of the probabilities of the cytoplasmic movers follow a uniform distribution, which is expected under the assumption of true null (Fig S6) (40). We used the standard Benjamini Hochberg multiple hypothesis correction procedure to assign q-values to the probability of movement (41). The Probable RNA polymerase II nuclear localization protein, SLC7A6OS, is one of our topmost hit with a median RNC shift of 0.53 and a q value of 7 e-9 (Table S1). But BACIQ is able to detect much more subtle changes. Among the newly assigned movers, BACIQ assigns highly significant q values of 1 e-4 to Nuclear export mediator factor, (NEMF) with 3.8% movement towards the nucleus. Even a change of subcellular localization as small as 1% results in a q-value of 0.025 for the 40S ribosomal protein S10, (RPS10) (Fig. 7C). With an estimated false discovery rate (FDR) of 5%, we detect ~750 putative Exportin-1 substrates, accounting for 612 unique gene symbols. Importantly, by reanalyzing previously published data with our newly developed statistical tool we discover ~2x more significant LMB responders (Exportin-1 substrates) as compared to our previous naïve analysis at an identical FDR (Fig. 7D). The Görlich and Chook groups identified Exportin-1 substrates in orthogonal approaches using either an affinity assay (42) or by curating a list from the literature (43). Our list of putative Exportin-1 substrates above the 5% FDR threshold exhibits a highly significant overlap with both these resources (p-values 5.5 e-29 and 3.1 e-6 respectively, hypergeometric test) (Fig. 7E). While there is still substantial disagreement between the three compared studies the overlap seems encouraging and might point towards an emerging consensus of Exportin-1 substrates. We believe the overlap is particularly meaningful due to the drastically different approaches by which these datasets were created.

## Discussion

We have shown how a hierarchical Beta-Binomial model can be used to adequately reflect uncertainty in quantitative multiplexed proteomics measurements. We presented the method and the implementation of a modeling pipeline which can assign confidence for both single peptide and multiple peptide proteomic measurements. We demonstrated how to estimate a calibration multiplier for a given instrument and mass resolution and then use that multiplier to convert a continuous MS-signal value into discrete event counts suitable for Beta-Binomial modeling. While we mostly demonstrate our approach in this paper for two condition scenarios, the entire framework can also be applied to multi condition cases. We validated our approach on the peptide and protein levels in synthetic samples for which we knew the true answer (Fig. 2, 6). To evaluate BACIQ in a real world scenario we re-analyzed a previously published dataset of subcellular localization movement upon inhibition of Exportin-1 with Leptomycin B. Without having to perform any additional experiments we increased the number of highly confident Exportin-1 substrates by ~2x. The mathematically rigorous integration of ion-statistics with concordance between different peptide measurements mapped to the same protein allowed us to confidently identify movement of some proteins by ~1% towards the nucleus as significant. The comparison with resources from other groups allowed us to independently validate this approach (42, 43) and points towards an emerging consensus on Exportin 1 substrates.

So far, we have only tested the BACIQ approach for data acquired with TMT-MS3 and TMTc+. We expect that with some adaptations BACIQ might be adequate for MS1–based labeled quantitative proteomics methods like SILAC or reductive methylation (44, 45). The systematic error associated with MS2-based multiplexed measurements will lead to inadequate confidence intervals for the underlying true protein ratios (11). Nevertheless, even when measurements are systematically distorted towards a 1:1 ratio due to interference, we suspect that BACIQ could still be useful for the prioritization of systematic changes in an experiment. In summary, we anticipate that our new statistical tool BACIQ will be highly valuable for researchers wanting to identify significant changes in proteomics studies and help to optimize the allocation of valuable resources for follow up studies.

## Acknowledgements

We would like to thank Graeme McAlister, Alexander Makarov, Yarden Katz and Josh Akey for helpful suggestions and discussions. Thanks to Elizabeth Van Itallie for comments on the manuscript. Bob Carpenter and the Stan team for technical support. L.P was supported by R01HD091846 and R01HD073104. This work was funded by the DOE Center for Advanced Bioenergy and Bioproducts Innovation (U.S. Department of Energy, Office of Science, Office of Biological and Environmental Research under Award Number DE-SC0018420). Any opinions, findings, and conclusions or recommendations expressed in this publication are those of the author(s) and do not necessarily reflect the views of the U.S. Department of Energy. This work was supported by NIH grant 1R35GM128813.

## Abbreviations

RNC: Relative Nuclear Concentration
FDR: False Discovery Rate
TMT: Tandem Mass Tag
TMT-MS3: MultiNotch MS3 analysis of TMT samples
TMTc+: Complement Reporter Ion Quantification Method
ANOVA: Analysis of variance
BACIQ: Bayesian Approach to Confidence Intervals for protein Quantitation
NEM: N-Ethylmaleimide
BCA: bicinchoninic acid
Lys-C: Lysil Endopeptidase
EPPS: 4-(2-Hydroxyethyl)-1-piperazinepropanesulfonic acid
RCF: Relative Centrifugational Force
LMB: Leptomycin B
Exp1: Exportin 1

