## Supplemental Information for "Bayesian Confidence Intervals for Multiplexed Proteomics Integrate lon-Statistics with Peptide Quantification Concordance"

### Supplementary Figures

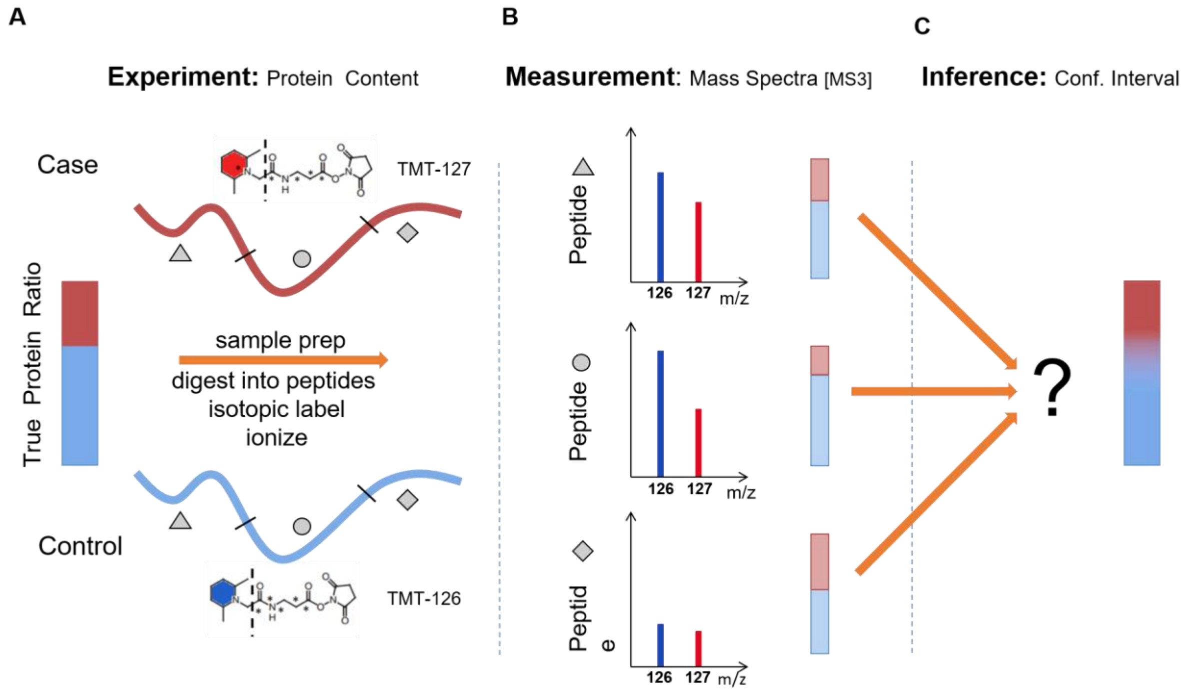


**Figure S1. Overview of the challenge to integrate multiplexed proteomics measurements into an estimation of protein ratio between conditions and the associated confidence in this measurement**. A) Multiplexed proteomics allows the comparison of protein abundance among multiple conditions. For simplicity, only two conditions and three peptide measurements are shown. A protein with true protein ratio (red-blue bar) is digested into peptides. The peptides are labeled with isobaric tags e.g. TMT to encode the different conditions. The digestion and labeling for different conditions is done separately. The peptides are combined, ionized, and injected into the mass spectrometer. Thus, the differential digestion and labeling could introduce disagreement in the peptide ratios. B) The different peptides derived from each protein results in separate spectra, which allow quantification using accurate multiplexed proteomics methods, e.g. MultiNotch MS3 or TMTc+. The relative intensity of the peptide signal can be used to quantify protein abundance. Each spectrum contains the information of relative abundance and signal i.e. the number of ions by which this ratio was measured. Thus, ion statistics introduce yet another source of distortion to the true ratio. C) The challenge is how the information of different peptide intensity and agreement/disagreement between measured peptide ratios can be integrated to accurately estimate the underlying true protein ratio with associated confidence.

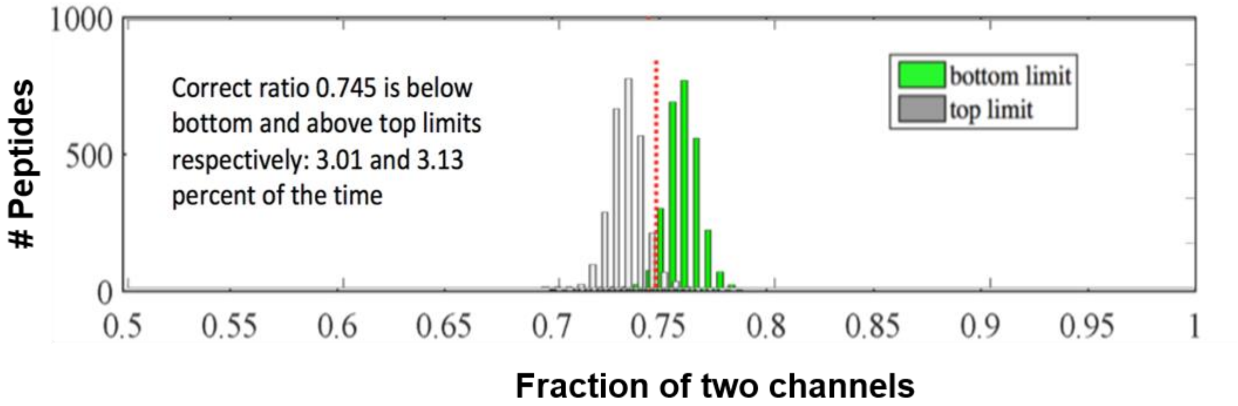


**Figure S2: Assignment of confidence intervals on the peptide level is adequate for various ratios.** This figure was constructed equivalently to Figure 2D, except the true ratio is about 3:4. The 95% confidence intervals are below bottom 3.01% and above top limits 3.13% of the time.

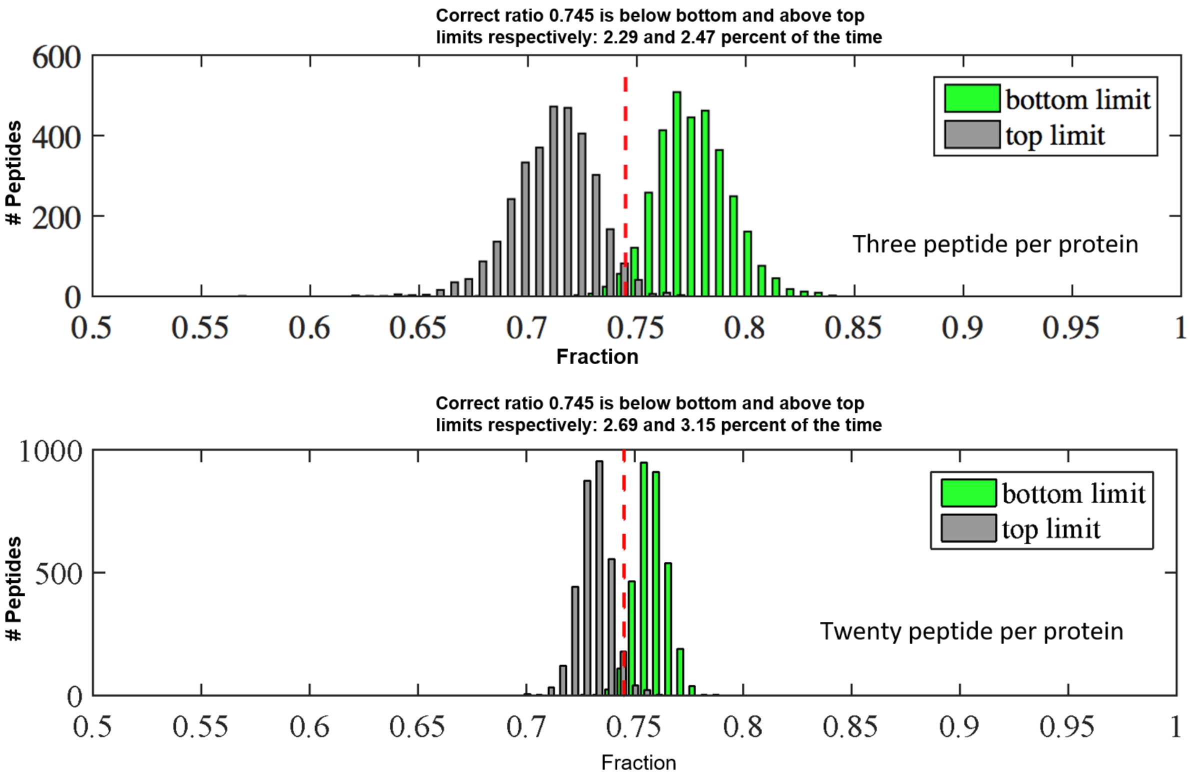


**Figure S3: Summing up peptide signal is adequate for the artificial protein case.** Artificial proteins are generated by summing from several peptides from a sample in which all peptide ratios are identical. A) 3 peptides were selected per artificial protein. The correct mixing ratio for this sample is 0.745. The 95% confidence intervals are below bottom 2.69% and above top limits 3.15% of the time. B) 20 peptides were assigned per artificial protein. Bottom limit of 95% confidence intervals is 2.29% above true ratio, and top limit is 2.47% of the time below true ratio.

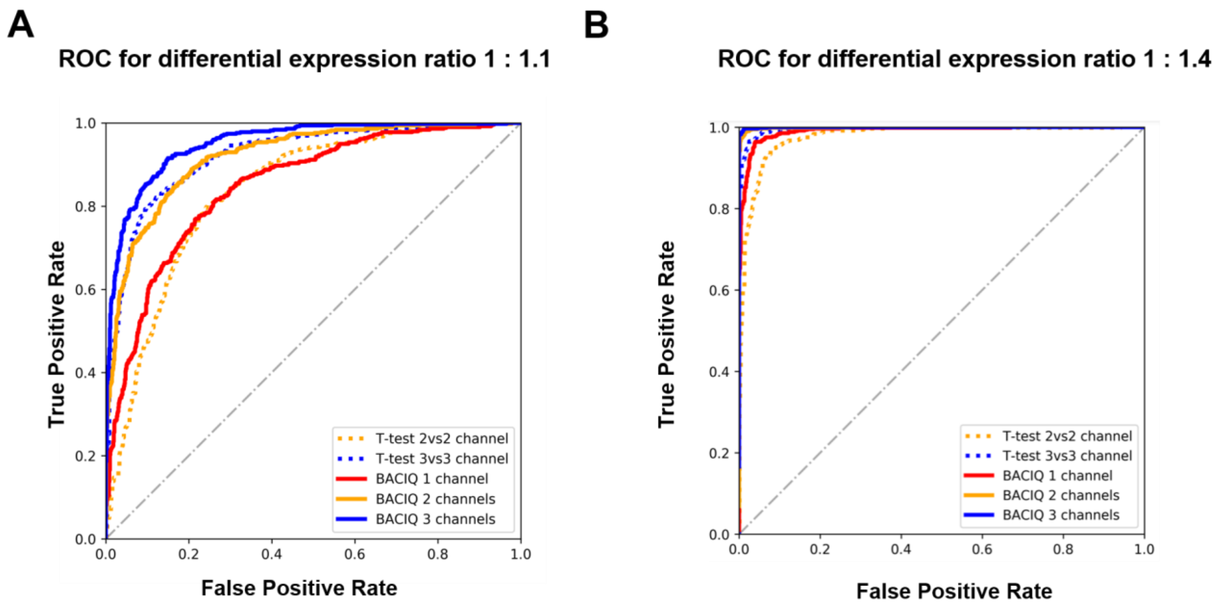


**Figure S4. Comparison of BACIQ with T-test for additional ratios of differentially expressed proteins.** This figure was equivalently constructed to Figure 5, but the ratio of the E. spiked in E. coli proteins was changed to A) 1:1.1 and B) 1:1.4. For all tested ratios BACIQ outperforms the naïve t-test in the ability to detect expression changes.

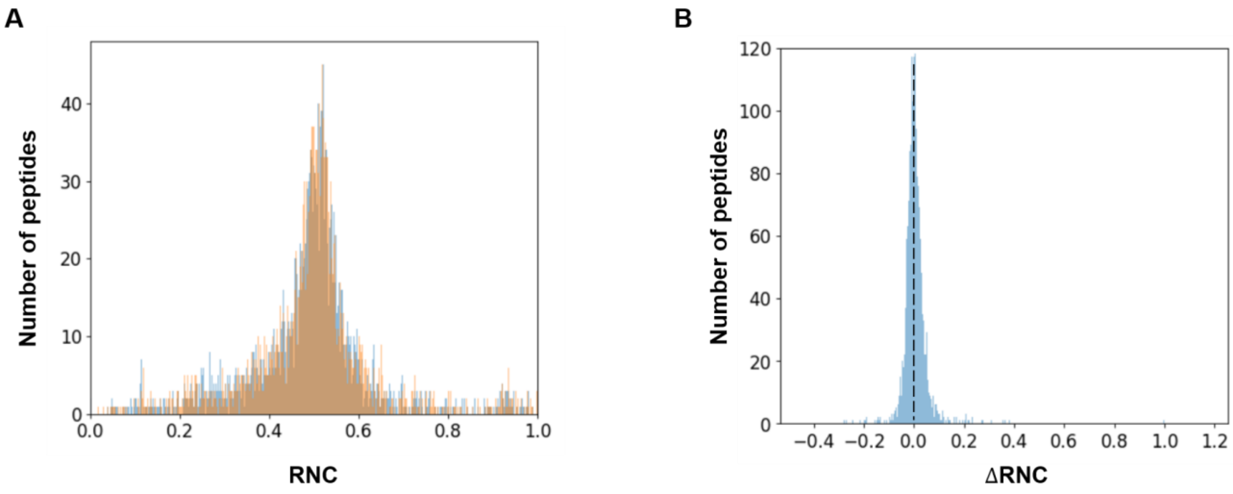


**Figure S5: The raw MS-signal is normalized according to the proteins that are small enough to diffuse and equilibrate via the nuclear pore.** A) The blue and the orange histogram represent RNC of the control and drug –treated samples for small proteins. B) The histogram of the normalized peptides from small proteins indicate a median RNC change of zero.

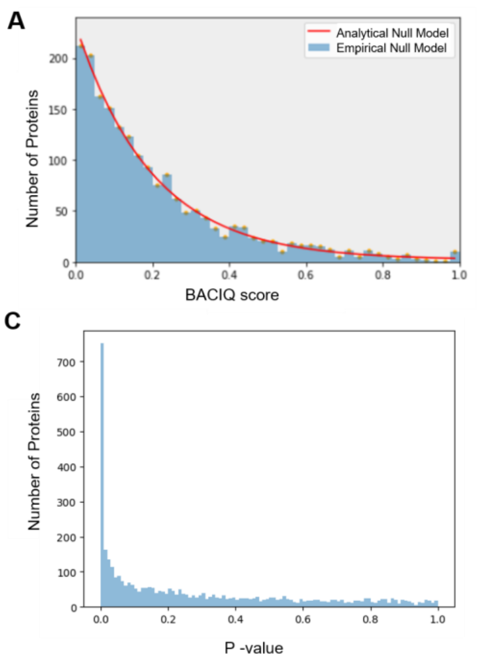


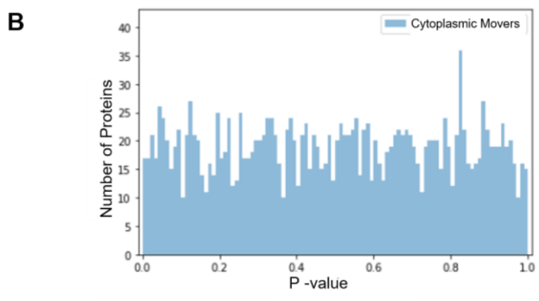


D

| Significance Level (down) | Bonferroni (FWER) | Benjamin-Hochberg (FDR) |
|---|---|---|
| | Total proteins | Total proteins |
| 1% | 160 | 454 |
| 5% | 200 | 757 |

**Figure S6: Multiple Hypothesis Corrections to assign FDRs to the shifters.** A) A histogram of the BACIQ scores of apparent cytoplasmic shifters. The red line is the analytic approximation. B) P-values associated with the BACIQ score of the cytoplasmic movers. The p values of the cytoplasmic shifters follow a uniform distribution indicating that the cytoplasmic shifters fulfill a key requirement for the true null. C) P- values of the nucleus movers. The peak close to 0 is where most of the alternative hypothesis lies along with some potential false positives D) Applying standard multiple hypothesis correction procedures (Bonferroni, or Benjamini-Hochberg) helps assign the False discovery rates to the probability values of the protein.

### Supplementary Material

#### Fitting the parameter to convert MS-signal to number of ions.

Number of successes y on n coin tosses, given its fairness p follow a binomial distribution $y \sim Bin(n,p)$. To functional form of the convergence of fraction of successes to the true fraction of successes (fairness of the coin) on increasing number of coin tosses n is given by the relationship of the CV of binomial distribution with n. The mean of the binomial distribution with parameters **n** (number of tosses) and **p** (probability of success) is $m = np$ and the standard deviation is $s = \sqrt{np(1-p)}$ thus the coefficient of variation is $Cv = \sqrt{\frac{1-p}{np}}$.

We fit a single parameter ***m*** as a multiplier to an S/N value ***s*** where ***n=ms*** to the binned data as illustrated in Figure 2B. The data of 10534 points is binned by 500 data points into 21 bins, and CV is calculated for each bin with MATLAB's Nonlinear-Least-Squares fitting method.

#### Comparison of BACIQ to t-test

We compare the t-test, which uses 2 or 3 repeats, to BACIQ. The t-test ranks the proteins from most to least likely differentially expressed, by 1-pvalue. The p-value is the probability that the different sets of peptide measurements for different conditions do not correspond to the differential expression, rather due to measurement noise. BACIQ ranks the proteins based on the probability mass falling to the right of 0.5 in the distribution for fraction between two conditions. Sliding threshold on the ranking allows building ROC curve – a way to compare probabilistic classifiers' tradeoff between true positive and false positive rates.

#### Implementation of BACIQ for the Exportin 1 inhibition data

BACIQ was first run on the nucleus and the cytoplasm channel to obtain the RNC distribution for the control and again on the nucleus and cytoplasm channel to obtain the RNC for drug treated sample. We considered the 7841 proteins that have more than one peptide. The samples for the marginal posterior distribution of the true protein ratio were collected by dividing the RNC space (0,1) into 40000 bins, giving the grid size of 2.5e-5 RNC.

#### Computing the probability values for the Exportin 1 inhibition data

To calculate the probability of shifting from the RNC distributions of control and drug treated samples, we ask the $P(RNC_{drug} > RNC_{control})$ for the proteins with positive shift RNC ( $shift\ RNC = Median\ RNC_{drug} - Median\ RNC_{control}$) and $P(RNC_{control} > RNC_{drug})$ for the negative shift RNC. These probabilities can be numerically computed as shown in the figure S6 or directly using the following formulae:-

$$P(RNC_{control} > RNC_{drug}) = \int_0^1 cdf_{drug}(RNC)\, pdf_{control}\,(RNC)\, d\, RNC$$
$$= \sum_1^{40000} cdf_{drug}(i)\, pdf_{control}\,(i)$$
$$= Area\ to\ the\ left\ of\ 0\ in\ shift\ RNC\ curve$$

$$P(RNC_{drug} > RNC_{control}) = \int_0^1 cdf_{control}(RNC)\, pdf_{drug}\,(RNC)\, d\, RNC$$
$$= \sum_1^{40000} cdf_{control}(i)\, pdf_{drug}\,(i)$$
$$= Area\ to\ the\ right\ of\ 0\ in\ shift\ RNC\ curve.$$

**Normalization and multiple hypothesis correction**

To correct for pipetting errors, we normalized corresponding nuclear and cytoplasmic proteins, so that the median of proteins in complexes smaller than 40kDA is distributed 1:1 (RNC = 0.5)(1) (Fig. S5).

To counteract detection of apparent LMB movers due to slight off normalization or isotopic impurities of the TMT tag, we additionally, normalized the LMB treated data so that there were equal number of cytoplasmic and nucleus movers for high t-test p value scores. Please note that this approach is conservative and will lead to lack of sensitivity but increase accuracy in identifying the true nucleus movers.

We used the distribution of the BACIQ probability scores for cytoplasmic movers (before overcorrection) as model for the null. To accurately calculate the P-value associated with the BACIQ scores for all the proteins, the empirical null model is approximated by an analytical model. The binned data was fit to the exponential distribution using Non-Linear Least Squares method (Fig. S6A). A key property of the null distribution is that its p-values are uniformly distributed. A uniform distribution of the p values for the BACIQ scores of cytoplasmic movers suggest that they indeed correspond to the true null (Fig. S6B). On the other hand, the p-values for the nucleus movers show a large peak close to 0 (Fig. S6C). Fig. S6D tabulates the number of proteins being pulled out as significant on applying the two standard but conservative multiple hypothesis correcting procedures

**Comparison of putative Exportin-1 substrates with other databases.**

We compared proteins we identified as significantly moving towards the nucleus to the Crm-1 binder from Xenopus published by the Görlich group (2) and a database curated from literature (NESDB)(3). To facilitate comparison of proteins identified with an mRNA derived reference database, we mapped all Xenopus proteins to their human gene symbols as previously described (4).

All the proteins that were called Cargo A, Cargo B, or low abundant Cargo by Görlich were considered as "binders" for the overlap. The 971 binder proteins collapsed down to 960 unique gene symbols. For our study, the 747 proteins at 5% FDR for our dataset collapsed down to 612 unique gene symbols. We found an overlap of 245 unique gene symbols between both the datasets of unique gene symbols. To calculate the statistical significance of this overlap, we calculated the probability of observing an intersection of at least 245 on overlapping 960 genes and 612 genes from the total of 5531 unique gene symbols identified in our data set, by random chance. The small p-values (5.5 e-29 ) using the hypergeometric distribution indicate that this overlap is significant. Similarly, for the 230 unique gene symbols of NESDB, only 76 were given any predictions from our dataset. Out of these 76, 26 overlapped with the 612 unique gene symbols from our study. To calculate the statistical significance of this overlap, we asked the probability of observing an intersection of at least 26 on overlapping 76 genes with 612 genes from the total of 5531 unique gene symbols. The small p-values (3.1 e-6) using the hypergeometric distribution indicate that this overlap is significant.
